# Cellular proximity and conditional coordination in the colorectal and pancreatic tumor microenvironment: a patient-level analysis of multiplexed imaging

**DOI:** 10.64898/2026.10.08.757632

**Authors:** Sagnik Bhadury, Timothy L. Frankel, Arvind Rao

**Affiliations:** Department of Epidemiology, Biostatistics and Informatics, University of Pennsylvania, Philadelphia, PA, USA; Department of Surgery, University of Michigan, Ann Arbor, MI, USA; Department of Computational Medicine and Bioinformatics, University of Michigan, Ann Arbor, MI, USA; Department of Biostatistics, University of Michigan, Ann Arbor, MI, USA; Department of Biomedical Engineering, University of Michigan, Ann Arbor, MI, USA

**Keywords:** tumor microenvironment, multiplexed imaging, spatial statistics, conditional association, cellular proximity, patient-level inference, colorectal cancer, pancreatic cancer

## Abstract

Immune cells in the tumor microenvironment are organized, and in colorectal and pancreatic cancer their arrangement around the tumor epithelium carries prognostic information. Multiplexed tissue imaging records the position and phenotype of every cell in a section, and studies of such images now routinely report that two cell types are spatially associated. However, the summaries behind these reports answer different questions. A proximity statistic asks whether two cell types lie close together more often than expected by chance. A conditional network asks whether their local densities rise and fall together once the other cell types are held fixed. The two can disagree, because every cell type is abundant where tissue is cell-dense and because the images of one patient are repeated measurements of that patient. We asked how epithelial and immune cells are organized in two settings: colorectal cancer with a Crohn’s-like reaction (CLR) or diffuse inflammatory infiltration (DII), in 140 CODEX images from 35 patients, and pancreatic ductal adenocarcinoma (PDAC) against its precursor, intraductal papillary mucinous neoplasm (IPMN), in 207 multiplex immunofluorescence images from 141 patients. We developed a patient-level analysis that keeps proximity, recurrent local composition and conditional coordination as separate quantities, tests conditional organization as a function of local epithelial density within each section, checks before every conditional estimate that the two cell types are disjoint sets of cells observed together in enough images, and calibrates each conclusion by patient-label permutation, cellularity adjustment under two definitions, spatial-scale perturbation and fully nested leave-one-patient-out prediction. In colorectal cancer, the conditional association between epithelial and myeloid density turned negative with local epithelial density about four times faster in DII than in CLR (difference *−*0.0553 per quartile; 95% CI, *−*0.0833 to *−*0.0273; patient-label familywise *P* = 0.006), with no accompanying difference in proximity. In the pancreas, cytotoxic and helper T cells were closer together in PDAC than in IPMN at every scale examined (*q ≤* 0.023), whereas their conditional association differed between the two diseases under only one of the two cellularity definitions. Recurrent neighborhoods carried the most patient-level information in colorectal cancer (AUC 0.794) and combined spatial features in the pancreas (AUC 0.771). Pancreatic epithelial–immune and regulatory T-cell quantities could not be estimated from these cores and are reported as such. The analysis is implemented in the R package spatialMIFtools, available at https://github.com/sagnikbhadury/spatialMIFtools.

## 1 Introduction

### Disease burden

Colorectal cancer and pancreatic cancer are the second and third leading causes of cancer death in the United States. In 2025 an estimated 154,270 new colorectal cancers and 52,900 deaths, and 67,440 new pancreatic cancers and 51,980 deaths, are expected, with five-year relative survival of 64% and 13% (Siegel et al., 2025). In both diseases the immune content of the tumor microenvironment carries prognostic information, and the question has moved from how many immune cells a tumor contains to how they are arranged with respect to the tumor epithelium and to each other. In colorectal cancer, a Crohn’s-like reaction (CLR), that is, discrete lymphoid aggregates at the tumor border, was described as a favorable host response three decades ago (Graham and Appelman, 1990). It is contrasted with diffuse inflammatory infiltration (DII), which lacks that organization and carries a worse prognosis (Schürch et al., 2020). In the pancreas, pancreatic ductal adenocarcinoma (PDAC) arises from precursor lesions that include intraductal papillary mucinous neoplasms (IPMNs), and the immune microenvironment of the invasive tumor differs from that of its precursors (Halbrook et al., 2023).

### Immune organization in the tumor microenvironment

What these observations share is coordination between cell types at a local scale. In CLR, the peritumoral aggregates mature toward tertiary lymphoid structures in which T cells and antigen-presenting cells sit together. Schürch et al. (2020) described coordinated tumor, T-cell and macrophage neighborhoods at the invasive front of CLR tumors and a more fragmented organization in DII. In PDAC, sub-microenvironments with distinct immune and fibroblast composition coexist within one section (Grünwald et al., 2021). Multiplex imaging has profiled epithelial cells, antigen-presenting cells, helper T cells, cytotoxic T lymphocytes (CTLs) and regulatory T cells across precursor lesions and invasive tumors (Enzler et al., 2024), and organized helper and cytotoxic T-cell neighborhoods have been associated with longer survival (Arseneau et al., 2025; Damle et al., 2026). Hence the biological hypothesis is not only that disease changes the abundance of immune cells. It is that disease changes whether the local densities of two cell types move together, and where in the section they do so.

### Multiplexed imaging and the two cohorts

Multiplexed protein imaging records the phenotype and location of every cell in a tissue section, and cohorts of such sections are now collected routinely in oncology (de Souza et al., 2024). We analyze two of them (Section 2). The first is the CODEX colorectal cancer cohort of Schürch et al. (2020): 35 patients (17 CLR and 18 DII), four tissue-microarray images per patient, 140 images and 258,385 phenotyped cells. The second is the multiplex-immunofluorescence pancreatic cohort that served as the source data of the published ISPAT application (Bhadury et al., 2026b): 141 patients (70 IPMN and 71 PDAC), 207 images and 274,332 cells, each carrying one of five mutually exclusive lineage labels. Figures 1 and 2 show one image per pathology from each cohort as a smoothed epithelial-density surface. Two properties are visible there. Epithelial density varies strongly within a single section, and its range differs between sections of the same cohort. Hence a question about organization has to be posed relative to the local tumor density of each section, and we build the analysis around that coordinate.

**Figure 1:**
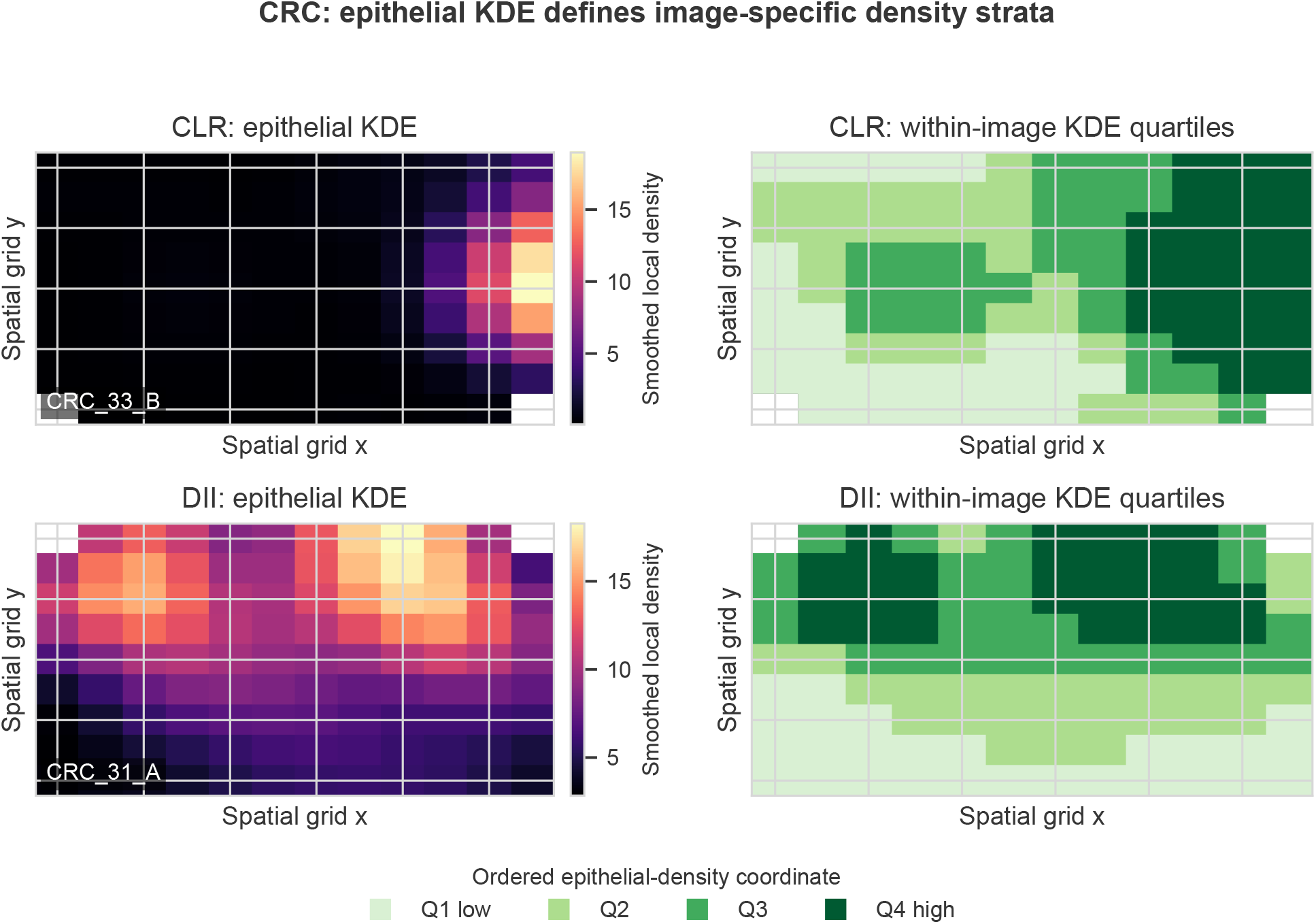
Construction of the CRC epithelial-density coordinate. Left, smoothed epithelial density for one CLR and one DII image. Right, the corresponding within-image quartile assignments; each quartile holds about one fourth of the occupied bins. Q1–Q4 preserve spatial ordering within a section and impose no common absolute-density threshold across images. The panels illustrate preprocessing and do not by themselves estimate a CLR–DII difference.

**Figure 2:**
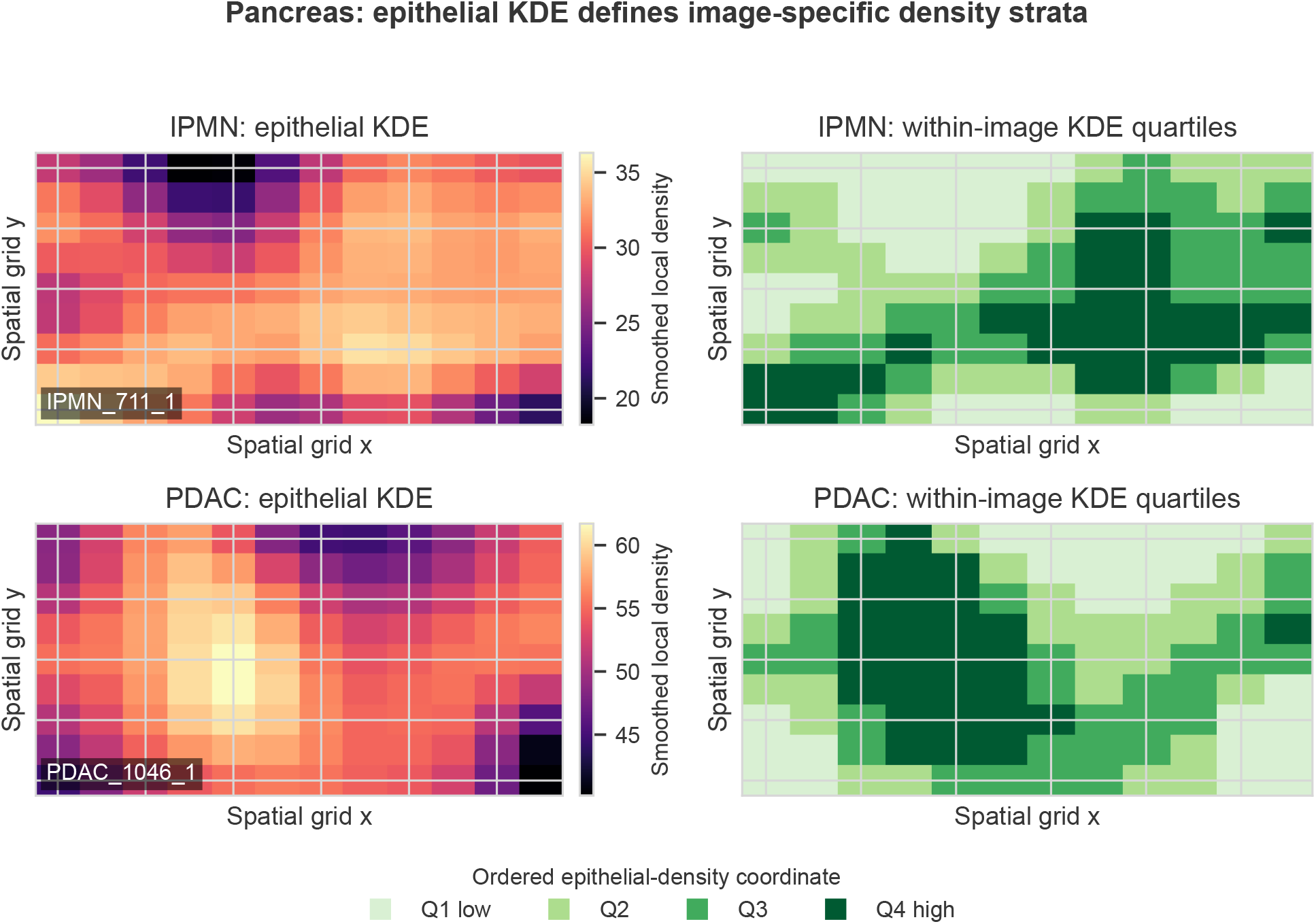
Construction of the pancreatic epithelial-density coordinate. The smoothed epithelial-density ranges differ between the displayed IPMN and PDAC images (left), whereas the quartiles encode relative position within each image (right). The rank transformation permits the same ordered analysis across sections with different density ranges. It neither equates their absolute epithelial densities nor represents temporal progression from IPMN to PDAC.

### Scientific question

We ask whether two cell types are organized with respect to each other in a disease-specific way. In colorectal cancer, we ask whether the association between epithelial and myeloid density changes with local epithelial density differently in DII than in CLR. In the pancreas, we ask whether cytotoxic and helper T cells are organized differently in PDAC than in IPMN, and whether that organization survives adjustment for local cell burden. In both cohorts, we ask how much patient-level information about pathology each spatial representation carries. Statistically, this translates to estimating three quantities from cell coordinates at the patient level, namely the excess of close cross-type cell pairs over chance, the prevalence of recurrent local mixtures, and the partial correlation between two density fields given the other cell types, and then testing whether each differs by pathology. Answering these questions raises four statistical challenges.

### Statistical challenges

First, the summaries in current use answer different questions. A proximity statistic conditions on nothing, a neighborhood model describes which local mixtures recur, and a conditional network holds the other measured cell types fixed. Nothing forces the three to agree. Second, every cell type is abundant where tissue is cell-dense, so a disease that changes cellularity can move every pairwise summary (proximity, neighborhood prevalence and partial correlation alike) at once. Recent benchmarking shows that apparent group differences in cell–cell interaction can be driven by abundance alone (Siddiqui et al., 2026; Soupir et al., 2025), and the definition of the cellularity nuisance is a choice that defines the estimand. Third, proximity and neighborhood summaries depend on the radius, grid or bandwidth at which they are computed (Bull et al., 2024; Soupir et al., 2025), and one patient contributes several images, so images are repeated measurements and not independent observations (Samorodnitsky and Wu, 2024). With 35 patients, large-sample clustered inference is used near the lower end of its comfort (Mancl and DeRouen, 2001). Fourth, a partial correlation between two density fields can be interpreted only if the fields are computed from disjoint sets of physical cells, and it can be estimated at the patient level only if enough images carry both cell types. In tissue-microarray cohorts some pairs of compartments rarely occupy the same core, and to our knowledge neither condition is checked in current practice.

### Existing analyses

Broadly, existing analyses of spatial organization in multiplexed imaging fall into three classes, which reviews of the field organize by the property of tissue they measure (de Souza et al., 2024; Lopez Janeiro et al., 2024). The first class measures marginal proximity: point-process and correlation-function statistics (e.g., Ripley’s cross-*K* and its extensions) that operate on intercellular distances and test co-localization against a spatial null (Barua et al., 2018; Bull et al., 2024; Soupir et al., 2025). However, these statistics condition on no other cell type, and without an abundance-appropriate reference they register tissue morphology instead of association (Siddiqui et al., 2026). The second class describes recurrent local composition: neighborhood models that cluster local mixtures, which were the basis of the CLR–DII distinction (Schürch et al., 2020), and learned graph-based or segmentation-free representations that carry patient-level information beyond composition (Wu et al., 2022; Baranwal et al., 2021; Tan et al., 2025). However, a neighborhood is a mixture and not a pairwise dependence, and a learned representation does not return an interpretable estimate for a named pair of cell types. The third class estimates conditional networks. ISPAT estimates shared and region-specific partial correlation structure across zones of a section (Bhadury et al., 2026b), GP-GHS estimates coefficients that vary continuously across a section (Bhadury et al., 2026a), and ISPAT-3D extends zone-specific networks to volumetric imaging (Bhadury and Rao, 2026). However, these are estimators for one image and were validated as such. None of them checks that the two compartments are separable and supported, adjusts for cellularity under more than one nuisance definition, or carries an image-level estimate to a patient-level inference with the repeated-measurement structure respected. Reporting standards for multiplex immunohistochemistry and immunofluorescence now ask for density adjustment, documented image selection, spatial-scale reporting and patient-level validation (Sater et al., 2025; Taube et al., 2025), but they do not say how these are to be done together. In summary, none of the three classes keeps marginal and conditional quantities apart under one patient-level standard, which is what the two properties visible in Figures 1 and 2 require. To address these challenges, we propose a novel patient-level analysis that separates cellular proximity from conditional coordination.

### Contributions

Our analysis makes four contributions, one per challenge. First, it treats marginal proximity, recurrent local composition and conditional coordination as distinct estimands, with the patient as the sampling unit and images and spatial bins as nested observations (Section 3.1). Second, it estimates cellularity-adjusted networks under two nuisance definitions, total cellularity and total cellularity with the focal counts excluded, so that a conditional conclusion is reported together with the definition it depends on (Section 3.5). Third, it introduces a within-section epithelial-density coordinate, defined by image-specific quartiles of a smoothed epithelial surface, along which conditional organization is tested as a function of local tumor density by a patient-clustered ordered model, calibrated by patient-label permutation and checked for spatial scale (Sections 3.3 and 3.6). Fourth, it checks before every conditional estimate that the two compartments are disjoint sets of cells observed together in enough images, and it reports quantities that fail this check as non-estimable and not as null (Section 3.1). Every conclusion is then subjected to the same checks: cellularity adjustment, spatial-scale and bandwidth perturbation, neighborhood resampling, patient-label permutation, comparison with two published estimators, and fully nested leave-one-patient-out prediction. Unlike a single-summary analysis, this analysis can report that proximity and coordination disagree. Unlike an image-level analysis, it never counts a bin or an image as a sampling unit. The main cost is that some quantities a single-summary analysis would report as null are reported here as non-estimable, and the pancreatic epithelial–immune and regulatory T-cell contrasts are such quantities.

Applied in parallel to the two cohorts, our analysis returns a different answer in each. In colorectal cancer, the conditional association between epithelial and myeloid density turns negative with local epithelial density faster in DII than in CLR, the contrast survives patient-label randomization, and proximity shows no corresponding difference. In the pancreas, cytotoxic and helper T cells are closer together in PDAC at every scale examined, whereas their conditional contrast is removed by adjustment for total cellularity and retained when the focal T-cell counts are excluded from the nuisance burden. The two cohorts are parallel applications of one standard and not replications of one biological result. ISPAT and GP-GHS appear only as comparators, because reanalysis of a cohort already used to develop a method is not independent validation of that method.

Two scope limits apply throughout. The pancreatic source annotation contains no positive regulatory T-cell calls, and pancreatic epithelial and immune compartments almost never share a core, so the corresponding quantities are non-estimable and are neither replaced by a proxy nor read as null effects. More generally, the panels do not measure antigen presentation, T-cell activation, cytotoxicity, ligand–receptor binding or change over time. In this paper, interaction refers only to a statistical interaction term, and edge, association and coordination refer to conditional statistical relationships (none denotes a causal or physical cell–cell interaction). We call the CRC CD68^+^/CD163^+^ compartment myeloid throughout, because the panel does not establish antigen-presenting function. Predictive analyses quantify patient-level discrimination between pathology groups, and we describe them as spatial discriminative signatures and not as biomarkers.

### Roadmap

The rest of the paper is organized as follows. Section 2 describes the two cohorts and the phenotype compartments. Section 3 defines the sampling hierarchy and the estimands, the separability and support check, the spatial summaries, the inferential models, the comparison methods and evaluation metrics, the nested prediction and the simulation design. Section 4 reports the operating characteristics of the analysis on synthetic data and then the findings by cohort and cell-type pair, with their sensitivity, estimator comparison and predictive information. Section 5 interprets what the findings mean for the study of the tumor microenvironment and where the two cohorts diverge. The Supplementary Material contains the permutation distributions, the cellularity-definition sensitivity table, the coordinate-normalization comparison and the simulation summaries. The analysis is implemented in the open-source R package spatialMIFtools, available at https://github.com/sagnikbhadury/spatialMIFtools.

## 2 Two multiplexed-imaging cohorts

### Colorectal cancer

The CRC cohort contained 258,385 phenotyped cells in 140 CODEX images from 35 patients (17 CLR and 18 DII; four tissue-microarray images per patient) (Schürch et al., 2020). The source provides one categorical cell-type label per cell. From it we formed a predefined five-compartment representation: tumor cells as Epithelial, CD68^+^ or CD163^+^ macrophages as the CRC myeloid compartment, regulatory T cells as Treg, CD8^+^ T cells as CTL, and CD4^+^ or memory CD4^+^ T cells as HelperT. No cell belongs to two compartments. The CLR and DII labels are patient-level pathology groups assigned by the source study, and we use them as given.

### Pancreas

The pancreatic analysis used the source dataset of the published ISPAT application (Bhadury et al., 2026b). It contains one row per physical cell with a single mutually exclusive base type assigned from lineage markers: epithelial (cytokeratin), CD163 macrophage (reported as APC), CD8 T cell (CTL), CD4 T cell (HelperT) and regulatory T cell (Treg). PD-L1 descriptor columns do not participate in assignment. The export contains 141 patients (70 IPMN and 71 PDAC), 207 images and 274,332 cells. It is smaller than the sample reported in the published ISPAT application, and all pancreatic results in this paper refer to this 70/71-patient export. The regulatory T-cell column is uniformly negative in it, so all regulatory T-cell quantities are non-estimable. IPMN and PDAC are independent groups of patients, so nothing in this paper estimates within-patient progression.

### Why exclusive assignment matters

Because no cell belongs to two compartments, every conditional edge and every cross-pair statistic relates two disjoint sets of physical cells, and a partial correlation cannot relate a count to a subset of itself. This property is what makes the conditional quantities of Section 3.5 interpretable at all, and we verify it for every pair before any estimate (Section 3.1).

### What the exports do not contain

No clinical covariate, outcome, acquisition batch or pixel-to-micron calibration was available in either export. Hence proximity is measured in fractions of a characteristic image length and not in micrometres (Section 3.7), and no analysis adjusts for treatment, stage or batch.

### The epithelial-density landscape

Figures 1 and 2 show one image per pathology from each cohort as a smoothed epithelial-density surface, together with the within-image quartiles that define the coordinate used throughout (Section 3.3). We chose the displayed images without reference to any edge or pathology estimate, by ranking on the number of estimable zones and then on the number of occupied bins. The panels explain the construction and do not by themselves test anything.

## 3 Statistical analysis

In this section we define the sampling hierarchy and the three estimands, the check that precedes every conditional estimate, the spatial summaries, the inferential models, the comparison methods and metrics, the nested prediction and the simulation design. Every quantity below is derived from the cell coordinates, the compartment labels and the patient-level pathology label, and the derivation never uses the pathology label until the final contrast. This ordering is what later makes patient-label permutation exact for the label null.

### 3.1 Sampling hierarchy, estimands, and the separability and support check

Suppose the data consist of *P* patients, with *P* = 35 in CRC and *P* = 141 in the pancreas. Let *p* = 1, …, *P* index patients and *i* = 1, …, *I_p_* index the images of patient *p*, with *I_p_* = 4 in CRC and *I_p_ ≥* 1 in the pancreas. For each image the observed datum consists of (i) the coordinates (*x_pic_, y_pic_*) of its cells *c* = 1, …, *C_pi_* in the original image coordinate system, (ii) one compartment label *f* (*c*) *∈ {*Epithelial, Myeloid or APC, Treg, CTL, HelperT*}* per cell, and (iii) the patient-level pathology label *G_p_ ∈ {*0, 1*}*, coded 1 for DII in CRC and for PDAC in the pancreas. Every other quantity in the paper (bins, densities, neighborhoods, edges and enrichments) is derived from these three objects.

We analyzed the two cohorts in parallel and never pooled them. The independent sampling unit was the patient. Images were repeated fields nested within patients, and spatial bins were repeated observations nested within images. Pathology contrasts are oriented as DII minus CLR in CRC and PDAC minus IPMN in the pancreas, and all tests were two-sided. Both cohorts were existing studies, so no prospective calculation fixed a minimum detectable effect. We report patient-level effect estimates with their uncertainty, and the number of bins is never the inferential sample size.

Three estimands are kept apart throughout. Marginal proximity is the excess of close cross-phenotype cell pairs over a random-label null, computed on the raw coordinates. Recurrent local composition is the prevalence of learned multivariate neighborhood profiles. Conditional coordination is the partial correlation between two phenotype density fields given the others. Nothing in the analysis requires these to agree, and the question is which of them carries a pathology contrast that survives the checks. Note that the three estimands also differ in what they condition on. Proximity conditions on nothing, composition on the joint profile, and coordination on every other measured field, so a disease that changes only shared cellularity can move the first without moving the third.

Unless stated otherwise, we controlled multiplicity by the Benjamini–Hochberg false-discovery rate (FDR) within cohort and within a predefined analysis domain, with *q <* 0.05 taken as significant. The rule sorts the *m P* values of a domain in increasing order and rejects the hypotheses up to the largest *k* with *P*_(_*_k_*_)_ *≤* 0.05 *k/m*, so that the expected proportion of false discoveries among the edges called is at most 5%. We chose FDR over familywise control for two reasons. First, each domain is a screen of a small number of prespecified pairs in which a missed edge and a falsely called edge are comparably costly for a descriptive study of organization. Second, the primary conclusions are in any case calibrated by patient-label permutation (Section 3.6) and not by the *q* value alone.

Before any conditional estimate, we distinguished compartment separability from inferential estimability. Compartment separability requires the two density fields to be derived from disjoint physical cells. Inferential estimability additionally requires each pathology group to contribute at least 10 patients and at least 10 images with nonconstant densities for both compartments. We fixed this rule without reference to any pathology contrast. Edges failing it could be computed in isolated images, but we did not interpret them as patient-level effects.

### 3.2 Spatial binning and density transformation

Let *n_pibf_* be the number of cells of phenotype *f* in bin *b* of image *i* from patient *p*, and let *A_ib_* be the bin area in the original image coordinate system. We divided each image bounding box into a primary 12 *×* 9 rectangular grid and excluded bins with fewer than five cells. The grid gives about one hundred bins per image. This is enough for a five-phenotype network in most images (Section 3.5), and it keeps each bin large enough that the five-cell rule removes only sparsely populated tissue. Both choices are varied in the scale sensitivity. Phenotype intensity was represented as

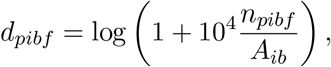

that is, a log cell count per fixed physical image area and not a within-bin proportion (the factor 10^4^ only fixes a convenient numerical scale). Bin-center coordinates were mapped separately to the unit square for spatial calculations, and this transformation did not enter *A_ib_*. The primary representation contained 12,081 occupied CRC bins and 12,244 occupied pancreatic bins.

For descriptive spatial structure, and as prediction features, we computed a four-nearest-neighbor auto-correlation score for each phenotype and image. With centered bin values *z_b_* = *d_b_ − d̄* and *N*_4_(*b*) the four nearest bin centers,

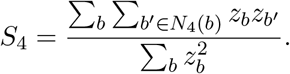

This score is proportional to Moran’s *I* on a regular four-neighbor graph but omits the normalization by total spatial weight, so we do not read its absolute magnitude as Moran’s *I*. A two-sided randomization *P* value used 199 permutations of the centered values within image. Images with fewer than 10 retained bins or a constant phenotype were non-estimable. Because patient summaries of *S*_4_ were standardized within the training fold before entering any predictive model, the omitted constant does not change the fitted predictor.

### 3.3 The epithelial-density coordinate

The biological question is posed relative to local tumor density, so we need a coordinate that places every bin of a section on a common scale of epithelial density without imposing one absolute threshold across sections. For each image, we placed the epithelial bin intensities on the grid and smoothed them with a Gaussian filter of standard deviation 1.25 grid bins, about one tenth of the horizontal extent. This width averages over adjacent bins without erasing the within-section gradient visible in Figures 1 and 2. We ranked the smoothed values at occupied bins within image and split them into four equally populated groups, Q1 to Q4 (ranking before quartiling makes assignments deterministic under ties). Hence Q1 and Q4 mark relatively low and high epithelial-density regions of the same section. They are neither absolute density thresholds nor stages of disease progression. Images with no varying epithelial surface were non-estimable.

Scale sensitivity used 8 *×* 6, 12 *×* 9 and 16 *×* 12 grids, with the Gaussian width scaled in proportion to horizontal resolution so that its image-scale support was retained, and, at the primary grid, standard deviations of 0.75, 1.25 and 2.0 bins. We compared neighborhood centers across grids after optimal one-to-one matching, disease-contrast edge ranks by Spearman correlation and sign agreement, and zone assignments by adjusted Rand index and exact agreement. The primary ordered-edge model was refitted at each bandwidth.

### 3.4 Recurrent spatial neighborhoods

Neighborhoods are recurrent multivariate bin-density profiles, and they do not alter the cell annotations. Within each cohort, we standardized the five phenotype intensities across retained bins and fit K-means with *K* = 3, …, 7 and 50 random starts to all bins. This range is the smallest that contains one profile per varying phenotype plus a low-density profile. We chose *K* by the largest silhouette coefficient on a fixed random sample of at most 6,000 bins. Names were assigned after fitting: a profile was named for its largest positive standardized coordinate among varying phenotypes, and profiles below the cohort mean on every varying coordinate were called Low-density. Naming did not affect assignments or inference. Patient-level prevalence was the proportion of a patient’s retained bins in each profile.

We evaluated stability in 200 stratified hierarchical bootstrap replicates. Within each pathology group, patients were sampled with replacement, then images with replacement within each selected patient, and replicates larger than 8,000 bins were subsampled to that size. The standardizer and K-means were refit with the frozen cohort-specific *K*, bootstrap centers were mapped to the reference scale and matched to reference centers by minimum total Euclidean distance (Hungarian algorithm), and a profile was called reproducible when its matched-center Pearson correlation was at least 0.8 in at least 80% of replicates.

### 3.5 Ridge conditional networks and cellularity adjustment

We estimated networks separately for every image with at least 12 retained bins. Let **Z***_pi_* be the *m_pi_ × r*matrix of standardized bin intensities for the *r* varying phenotypes in image *i*. We formed

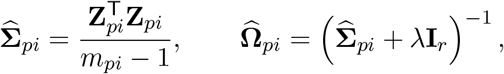

with *λ* = 0.15 in the primary analysis, and summarized the edge between phenotypes *f* and *g* by the ridge partial correlation

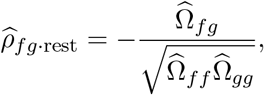

where “rest” denotes conditioning on all other varying density fields. A positive edge is a positive conditional association between density fields. It establishes neither contact, nor direction, nor signaling. Constant phenotypes, including pancreatic Treg, were dropped image by image. Each image costs one *r × r* inverse with *r ≤* 5, so the 24,325 bins of the two cohorts are processed in seconds, and the permutation and bootstrap loops dominate computing time (Section 3.12).

The ridge penalty has one job. With as few as 12 bins and up to five phenotypes, the sample covariance can be nearly singular, and its unpenalized inverse can return partial correlations near *±*1 from a handful of bins. Adding *λ* to the diagonal bounds the eigenvalues of the inverse away from infinity and shrinks every partial correlation toward zero, more strongly in images whose covariance is poorly conditioned. In other words, it trades a bias toward zero that is common to both pathology groups for a variance that no longer depends on a few bins. We fixed the value 0.15 before any pathology contrast was examined and refit the primary pancreatic conclusion at *λ ∈ {*0.05, 0.10, 0.15, 0.25, 0.50*}* (Supplementary Material).

For global pathology comparisons, we first averaged image edges within patient. The estimand was the difference between group means of these patient summaries, tested by Welch’s *t* test. In CRC, FDR was applied over the ten inferentially estimable edges. In the pancreas, the target-edge patient-label *P* value was primary, and BH values over the five computationally obtainable edges were kept only as a conservative sensitivity summary. Percentile 95% intervals used 2,000 stratified patient bootstrap replicates, and a hierarchical bootstrap that also resampled images within patient checked sensitivity to unequal image replication.

Because all intensities can rise together in cell-rich regions, we also estimated cellularity-adjusted networks. Within each image and phenotype we fit

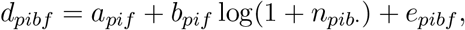

where *n_pib·_* is the total number of cells in the bin, and refit the ridge network to the residual matrix **E***_pi_*. This adjustment asks whether an edge persists beyond a linear association with local total cellularity. It does not adjust for unmeasured composition or technical batch.

The two nuisance definitions play different roles, and we report both because they answer different questions. Total cellularity, *n_pib·_*, removes everything that rises with local cell burden, including the two focal counts themselves. An edge that survives it is an association beyond any linear dependence on how crowded the bin is. However, when the two focal phenotypes make up most of the cells in a bin, the total is nearly a function of the focal counts, and residualizing on it removes the very variation the edge is computed from. The focal-excluded burden, *n*_other_ = *n*_total_ *− n*_CTL_ *− n*_HelperT_, used for the pancreatic CTL–HelperT edge, removes only the burden of cells outside the pair. An edge that survives it is an association beyond the non-focal crowding of the bin, but any shared dependence on the pair’s own abundance is left in place. Neither is the correct adjustment in general. Each defines an estimand, and when the two disagree we report the disagreement. We refit unadjusted, total-cell-adjusted and focal-excluded networks at the five ridge penalties. Among bins of images with nonconstant CTL and HelperT densities, we summarized the median, interquartile range and zero fraction of *n*_other_ and its Pearson correlation with total cellularity, which is the diagnostic that says whether the two definitions can differ.

### 3.6 Ordered inference along the coordinate and patient-label permutation

We estimated zone-specific ridge networks within each image and zone, requiring at least eight retained bins per image-zone, and formed analogous image-zone summaries for mean phenotype intensity and neighborhood prevalence. For each estimable edge, phenotype or neighborhood we fit the Gaussian generalized estimating equation

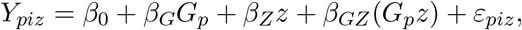

with *z ∈ {*1, 2, 3, 4*}* the ordered zone score and *G_p_* = 1 for DII or PDAC, repeated image-zone observations clustered by patient under an exchangeable working correlation, and robust sandwich standard errors. The parameters have the following interpretations. *β*_0_ + *β_Z_z* is the expected edge (or density, or prevalence) in the reference pathology, CLR or IPMN, at quartile *z*. *β_Z_* is its change per quartile of local epithelial density in the reference pathology. *β_G_* is the pathology difference extrapolated to *z* = 0 and is not interpreted. *β_GZ_*, the primary estimand, is the difference between pathologies in the change per epithelial-density quartile, so that *β_Z_* + *β_GZ_* is the change per quartile in DII or PDAC. Its Wald interval was 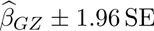, with FDR applied separately to edges, densities and prevalences within cohort. The linear zone score serves two purposes. It replaces four zone-specific contrasts by one parameter, and it gives the estimand a unit, a change per quartile, that the biological question is stated in. Its cost is that a non-monotone pattern across zones would be averaged away. As a secondary check, we computed patient-specific slopes after averaging images within patient and zone, for patients with at least three estimable zones, and compared them by Welch’s test.

With 35 CRC patients, the sandwich variance is used near the lower end of the cluster counts for which its large-sample approximation is comfortable, so we did not treat the Wald *q* value as sufficient by itself (Mancl and DeRouen, 2001). We therefore permuted pathology labels 999 times at the patient level within each cohort, holding the observed images, zones, densities and network estimates fixed. This conditioning is exact for the label null because none of those steps uses pathology (Section 3.1). Each CRC permutation refit all ten ordered edge interactions by GEE and repeated BH correction. Each pancreatic permutation refit the five computationally obtainable cellularity-adjusted contrasts by Welch’s test and repeated BH correction. All statistics were studentized, so that edges with different variances are compared on one scale. For the primary edge we computed a two-sided empirical *P* value from its absolute permuted statistic, and a familywise *P* value from the per-permutation maximum absolute statistic across the ten estimable CRC edges or, as a conservative sensitivity analysis only, across the five obtainable pancreatic edges. We also recorded the proportion of permuted datasets yielding any BH-adjusted *q <* 0.05.

### 3.7 Multiradius random-label cross-pair enrichment

We evaluated marginal proximity for predefined epithelial–immune and T-cell pairs on the original cell coordinates. No pixel-to-micron calibration was available for either cohort, so we rescaled each image by the single isotropic factor 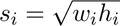, where *w_i_* and *h_i_* are its coordinate extents. This rescaling preserves aspect ratio and Euclidean distance up to one common factor, so a disc of radius *r* remains a disc. Hence the radii *r ∈ {*0.025, 0.05, 0.10*}* are fractions of a characteristic image length, and the resulting quantity is a dimensionless relative-distance enrichment and not a physical interaction radius. At each radius we counted unordered cross-phenotype cell pairs within distance *r*. Ninety-nine independent random relabelings of each binary phenotype vector preserved its number of positive cells while breaking spatial correspondence. For observed count *O_if_* (*r*) and mean permuted count 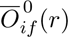

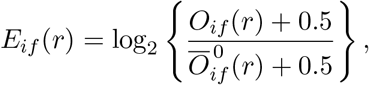

with the 0.5 preventing undefined ratios. Image enrichments were averaged within patient, groups were compared by Welch’s test, and FDR was controlled across pair–radius combinations within cohort. We call this quantity random-label cross-pair enrichment. It lacks the intensity normalization and boundary correction of Ripley’s cross-*K*. An earlier version scaled *x* and *y* separately, which maps every section to the unit square and alters its aspect ratio, so we computed and compared both normalizations. Aspect ratios were modest (CRC median 1.31, maximum 1.44; pancreas median 1.35, maximum 2.18) and no conclusion changed (Supplementary Material).

### 3.8 Comparison methods and cross-method concordance

Three network estimators appear in the paper, and we characterize each by what it estimates, how it was tuned and where it is weak. The ridge partial correlation of Section 3.5 is the primary estimator. It is closed-form, it is computable in every image with 12 bins, and its single tuning constant was fixed in advance. Its weakness is that it is a section-level or zone-level average and cannot locate where within a section an association is strongest. ISPAT (Bhadury et al., 2026b) estimates a shared network and zone-specific deviations across the Q1–Q4 zones of one image by variational Bayes spatial fitting and coordinate-ascent factor fitting. We used a Matérn kernel and an edge threshold of 0.15, the settings of the published application. Its weakness for the present purpose is that it is an estimator for one image and requires the zone labels as input, so it inherits the coordinate instead of testing it. GP-GHS (Bhadury et al., 2026a) estimates coefficients that vary continuously across a section under a graphical horseshoe prior. We used four basis functions, Matérn smoothness *ν* = 1.5, 800 MCMC iterations with 300 burn-in and thinning by five, an AND symmetrization rule and threshold 0.15, and the coefficient maps use the plotting convention that selects the nodewise direction with the smaller posterior shrinkage statistic. Its weakness is cost and mixing. All GP-GHS quantities in this paper come from these same 800-iteration fits, and targeted four-chain diagnostics on a representative CRC edge found a DII functional marginal against a strict *R̂* threshold and a strongly shrunk CLR functional with a mixing concern. For this reason GP-GHS is a comparator and not the basis of any primary statement. Both published methods were called through the Spatial Methods Workbench, an R Shiny environment that wraps them with their citations and settings (Bhadury, 2026).

We compared the published ISPAT and GP-GHS engines in an exploratory, availability-constrained CRC subset of 16 images, eight CLR and eight DII, with at most one image per patient, selected by number of estimable zones and then by retained-bin count. Previously fitted pancreatic models used a superseded overlapping-marker representation and were excluded from every cross-method comparison here. Disease contrasts were computed on identical successfully fitted image sets for each pair of methods. We summarized concordance by edge-direction agreement and by the Spearman correlation of absolute contrast ranks when at least five edges were estimable. GP-GHS maps were compared with Gaussian-localized ridge fields at the same grid locations by location-wise Spearman correlation, overlap of the top quartile of absolute edge magnitude, and agreement of the quadrant holding the strongest value. ISPAT and ridge were compared across the four matched zones by zone-wise rank, sign and strongest-zone agreement. These summaries quantify agreement in direction, magnitude rank and location. The method papers contain the validation of ISPAT and GP-GHS themselves.

### 3.9 Fully nested patient-level prediction and calibration

Four *L*_2_-penalized logistic models predicted DII in CRC and PDAC in the pancreas from (i) patient mean phenotype intensities (composition), (ii) neighborhood prevalences plus *S*_4_ scores, (iii) patient means and between-image standard deviations of all estimable ridge edges (network), or (iv) their combination. We fixed the inverse penalty *C* = 0.25 in advance instead of tuning it within fold, so that the nested pipeline learns features and not the penalty from 34 training patients, and inverse-frequency class weights kept the two pathology groups on equal footing in the loss.

Evaluation was leave-one-patient-out (LOPO). Within every fold, density standardization, selection of *K ∈ {*3, …, 7*}*, K-means centers, neighborhood assignments, median imputation and predictor standardization were learned from training patients only. K-means used 10 starts on at most 4,000 training bins with the silhouette on at most 2,500 sampled training bins. Held-out bins were assigned to the training centers, converted to prevalences and passed once through the fitted model. Features constant in a training fold were excluded. A held-out patient’s bins entered only that patient’s test-fold predictors, never the standardizer, clusters, imputer or regression. We recorded the selected *K* in every fold and the number of candidate predictors after excluding fold-constant features (this raw count is distinct from effective degrees of freedom). All assessment was internal, with no clinical covariates and no external cohort, so these analyses quantify discrimination and calibration of spatial representations and do not establish a deployable diagnostic.

### 3.10 Evaluation metrics

Every metric is defined here at the scale on which it is read. The ridge partial correlation 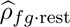 lies in [*−*1, 1]. Positive values mean that the two density fields rise together after the other fields are held fixed, and a pathology contrast is a difference of patient means of this quantity. The random-label enrichment *E_if_* (*r*) is zero when close cross-phenotype pairs are as frequent as under random labeling, positive when the two phenotypes are closer than that, and one unit is a doubling. A pathology contrast is again a difference of patient means. The silhouette coefficient, in [*−*1, 1], is the mean over bins of the difference between the distance to the nearest other cluster and the distance within the own cluster, divided by the larger of the two. Higher is better separated, and we used it to choose *K* because it does not require the clusters to be spherical or equally sized. Neighborhood reproducibility is the Pearson correlation between a bootstrap profile center and its matched reference center, in [*−*1, 1], summarized as the proportion of replicates at or above 0.8. The adjusted Rand index, in [0, 1] up to chance correction, measures agreement between two partitions, with 0 for chance agreement and 1 for identical partitions. We use it for zone assignments across bandwidths and for latent-profile recovery in simulation. Spearman’s rank correlation, in [*−*1, 1], measures agreement in ordering. We use it for edge ranks across grids, for absolute contrast ranks across estimators, and for GP-GHS against ridge fields across locations, because none of these comparisons is on a common scale.

Discrimination is the area under the receiver-operating characteristic curve (AUC), which is the probability that a randomly chosen DII or PDAC patient receives a higher out-of-fold score than a randomly chosen CLR or IPMN patient. It ranges from 0.5 for no discrimination to 1, and we chose it because it is threshold-free. Its 95% interval is the percentile interval from 2,000 class-stratified bootstrap samples, and its permutation *P* value comes from 999 permutations of the class labels against the fixed out-of-fold scores. Note that the latter tests association of the completed LOPO score with pathology and does not repeat model selection under each permutation. Calibration is the Brier score, the mean squared difference between the predicted probability and the 0/1 outcome, in [0, 1] with lower better and 0.25 for a constant prediction of one half at balanced prevalence, and the recalibration model logit {Pr (*Y* = 1)} = *a* + *b*logit (*p̂*), with ideal *a* = 0 and *b* = 1 and *b <* 1 indicating predictions that are more extreme than the outcomes warrant. Brier intervals used 2,000 class-stratified bootstrap samples, and calibration curves used four to ten quantile bins by cohort size. In simulation, a rejection rate is the proportion of replicates in which the target passes the FDR criterion, coverage is the proportion in which the 95% interval contains the generating value, and both carry a Monte Carlo standard error of 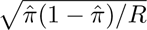 over *R*replicates.

### 3.11 Simulation experiments

The simulations test the components added for this application. They do not repeat the method-development experiments of ISPAT and GP-GHS (Bhadury et al., 2026b,a). Each module changes one part of the data-generating mechanism, and each contains a null cell so that the size of the procedure is measured beside its power. Where a dimension could be matched to the cohorts, we matched it.

#### Cellularity adjustment

We generated five smoothed Gaussian density fields for 40 patients, two images per patient and 81 bins per image. In one setting the two target phenotypes loaded more strongly on a common cellularity field in group 2 but had no residual edge. In a second, group 2 also had a residual association between the target fields. The observed cellularity covariate was a noisy version of the generating field, so the adjustment was deliberately misspecified in the direction the real data impose, where the measured burden is a proxy for the latent one. Ridge networks were estimated with and without residualization, all ten edges were compared between groups at the patient level, and BH correction was applied at 5%.

#### Ordered-zone inference

We generated data for 35 patients, four images per patient, four ordered zones and ten outcomes, with patient and image random intercepts inducing within-patient correlation. These are the CRC dimensions. Nine outcomes had zero group-by-zone interaction, and the target interaction was zero or *−*0.06. We chose this value to match the magnitude of the observed CRC estimate, so that the reported power refers to an effect of the size actually found. Each replicate used the exchangeable patient-clustered GEE of the data analysis, and we recorded the FDR-adjusted target rejection rate, false discoveries, bias and 95% interval coverage.

#### Cross-pair enrichment

We assigned two mutually exclusive phenotypes among 60 cells in each of two images from 30 patients, both spatially uniform under the null. Under the attraction alternative, 10 of the 15 group-2 cells of the second phenotype were placed with Gaussian displacement around randomly chosen cells of the first. Enrichment was computed at *r* = 0.05 with 99 relabelings, averaged within patient and compared by Welch’s test.

#### Nested prediction

We generated four latent neighborhood profiles in five dimensions for 30 patients with 40 bins each, with opposing pairs of profiles chosen to give the same expected composition. Under the alternative, group membership changed profile prevalence. Under the null it did not. Every LOPO training fold re-estimated standardization, selected *K ∈ {*3, 4, 5*}*, fit K-means, assigned held-out bins and fit the logistic model. We compared AUC from composition and neighborhood features and measured recovery of the latent profiles by adjusted Rand index and center root-mean-square error.

The network and GEE experiments used 500 replicates per setting, the cross-pair experiment 250, and the nested-prediction experiment 200. Monte Carlo standard errors were computed for rejection rates and mean AUCs.

### 3.12 Reproducibility and computing

Python-side stochastic stages used project seed 20260803 with deterministic offsets by analysis stage or held-out patient. Source paths, input checksums, parameters, serialized fits, fit status and derived-table locations were recorded in project manifests and run logs. Main analyses were implemented in Python 3.12. ISPAT and GP-GHS were called through the Spatial Methods Workbench, version 0.3.1 (Bhadury, 2026). On one workstation with an Intel Core i7-14700 processor and 32 GB of memory, the pancreatic primary pipeline from cell table to neighborhoods, ridge networks and leave-one-patient-out prediction ran in under six minutes. The 999 patient-label permutations of the unadjusted and adjusted pancreatic networks together with the fully nested 135-fold prediction took about two and a half minutes, the cross-pair enrichment for all 347 images under both normalizations with 99 relabelings each took just over one minute, and each 800-iteration GP-GHS fit took 0.4 to 3.4 seconds per image. The reusable implementation is the R package spatialMIFtools, with an optional Python backend through reticulate. Raw cell-level files were not modified or copied into the manuscript directory.

## 4 Results

### 4.1 Operating characteristics on synthetic data

Before the cohorts, we checked that each component of the analysis does what it is asked to do under its stated mechanism. When the two target phenotypes shared only a stronger cellularity loading, the unadjusted analysis selected the target edge in all 500 replicates, whereas the cellularity-adjusted analysis selected it in 0.008 of replicates (Monte Carlo SE, 0.0040) with a mean estimate of 0.0188. When a residual edge was generated, both analyses selected it in every replicate, and the mean adjusted estimate was 0.339. Hence the adjustment removes the specified shared-cellularity effect and keeps a direct edge, even though the observed covariate was only a noisy proxy for the generating field. For the ordered GEE at the CRC dimensions, the FDR-adjusted target rejection rate was 0.012 under the null and 0.898 when the interaction was *−*0.06, with a mean estimate of *−*0.0600 and 95% coverage of 0.958 under the alternative. Null coverage was 0.928, below nominal, which is the finite-cluster behavior of the unmodified sandwich interval at 35 patient clusters. This result is the reason the observed CRC statistic is calibrated by patient-label permutation and not by the Wald interval alone. The random-label cross-pair test rejected in 0.048 of null replicates (Monte Carlo SE, 0.0135) and in all replicates under the deliberately strong attraction alternative. Under the prediction null, mean AUC was 0.467 for composition and 0.492 for neighborhoods (Monte Carlo SEs, 0.0111 and 0.0087), and under the prevalence alternative composition AUC was 0.457 and neighborhood AUC was 1.000 (the latter reflects an intentionally well-separated alternative and is not an expectation for tissue data). In summary, the nested pipeline neither leaks nor manufactures signal, the ordered GEE has the power to detect an effect of the size found in CRC but under-covers at 35 clusters, and the cellularity adjustment behaves as intended under its stated mechanism. The graphical summaries and the latent-profile recovery statistics are in the Supplementary Material.

### 4.2 Analysis sets and which quantities could be estimated

The CRC analysis retained 12,081 grid bins from 140 images and 35 patients. The pancreatic cohort requires an explicit accounting, because retention rules remove images and therefore patients. Table 1 gives the attrition. Of 141 patients and 207 images, 135 patients and 191 images contributed at least one bin meeting the minimum-cell criterion, yielding 12,244 bins, and 121 patients contributed at least one image with enough bins for a conditional network.

**Table 1:** Pancreatic cohort provenance and analysis-set attrition. Each stage applies the retention rule stated in Section 3. No patient or image was removed on the basis of an estimated value.

| Stage | Patients |  | Images |  |
| --- | --- | --- | --- | --- |
|  | IPMN | PDAC | IPMN | PDAC |
| Source cells | 70 | 71 | 89 | 118 |
| Retained bins ($\geq 5$ cells/bin) | 70 | 65 | 89 | 102 |
| Network analysis ($\geq 12$ bins/image) | 66 | 55 | 82 | 83 |
| CTL–HelperT proximity ($\geq 3$ cells/type) | 33 | 45 | 37 | 56 |

Because epithelial values are ranked within image, an increase in epithelial density from Q1 to Q4 holds by construction, and what differs by pathology is its size. In CRC the DII-minus-CLR difference in linear change per quartile was 0.1289 (*q* = 0.00269), that is, a steeper within-section epithelial gradient in DII. These coefficients compare relative gradients within sections and are not estimates of change over disease time.

The separability and support check then determined which conditional edges could be interpreted. All ten CRC pairs were compartment-separable and inferentially estimable, with 127 to 139 contributing images each and no cell in two compartments. The exclusive pancreatic annotation made every evaluated pair compartment-separable, but only CTL–HelperT was inferentially estimable. The reason is structural. Of 207 pancreatic images, 99 contain at least three epithelial cells and 93 contain at least three CTLs, but only 5 contain both, and 6 contain at least three epithelial cells together with at least three helper T cells. The median per-image epithelial count is zero. In practice, cores are either epithelium-dominated or immune-dominated. Hence we report pancreatic epithelial–immune quantities as non-estimable and not as null results, and the same holds for all regulatory T-cell quantities.

### 4.3 Colorectal cancer: epithelial–myeloid coordination along local tumor density

In CRC, the primary quantity was the Epithelial–Myeloid partial correlation and its change across the epithelial-density coordinate. We find that this conditional association becomes more negative with local epithelial density in DII than in CLR. The DII-minus-CLR difference in the Q1–Q4 linear slope was *−*0.0553 per quartile (SE, 0.0143; 95% CI, *−*0.0833 to *−*0.0273). In interpretable units, the patient-averaged edge became more negative by 0.018 per quartile in CLR and by 0.072 per quartile in DII. In other words, as local epithelial density increases, the conditional association between epithelial and myeloid density fields turns negative about four times faster in DII. Patient-label randomization is the primary inferential statement for this result. Permuting pathology labels across patients 999 times gave a two-sided empirical *P* = 0.002 for the studentized target statistic, and *P* = 0.006 against the per-permutation maximum absolute statistic over all ten estimable edges. The clustered-GEE Wald test gave *q* = 0.00111 and is the secondary, model-based summary.

Two further analyses agreed, and two checks addressed artefacts of the construction. Estimating a slope per patient after averaging images gave a between-group difference of *−*0.0537. Refitting the interaction with the smoothing bandwidth changed from 1.25 bins to 0.75 and 2.0 bins met the FDR criterion at every setting (*q* = 0.0176, 0.0111 and 0.00580), so the result does not depend on the smoothing that defines the coordinate. Because DII sections are also more cell-dense along the coordinate, we checked whether the edge is a cellularity artefact. After residualizing each density on local total cellularity, the section-averaged Epithelial–Myeloid partial correlation remained negative in both groups, *−*0.202 in CLR and *−*0.271 in DII (difference *−*0.069; *q* = 0.57), in the same direction as the unadjusted section average, and 8 of the 10 CRC edge contrasts kept their sign under adjustment. Marginal proximity gave no corresponding signal. Under isotropic normalization, none of the three predefined CRC pairs differed between CLR and DII at any radius (all *q ≥* 0.45). Section-averaged conditional contrasts likewise gave no CRC edge meeting the FDR criterion (Figure 3A), which is expected, because a section average cannot show a change along the coordinate.

**Figure 3:**
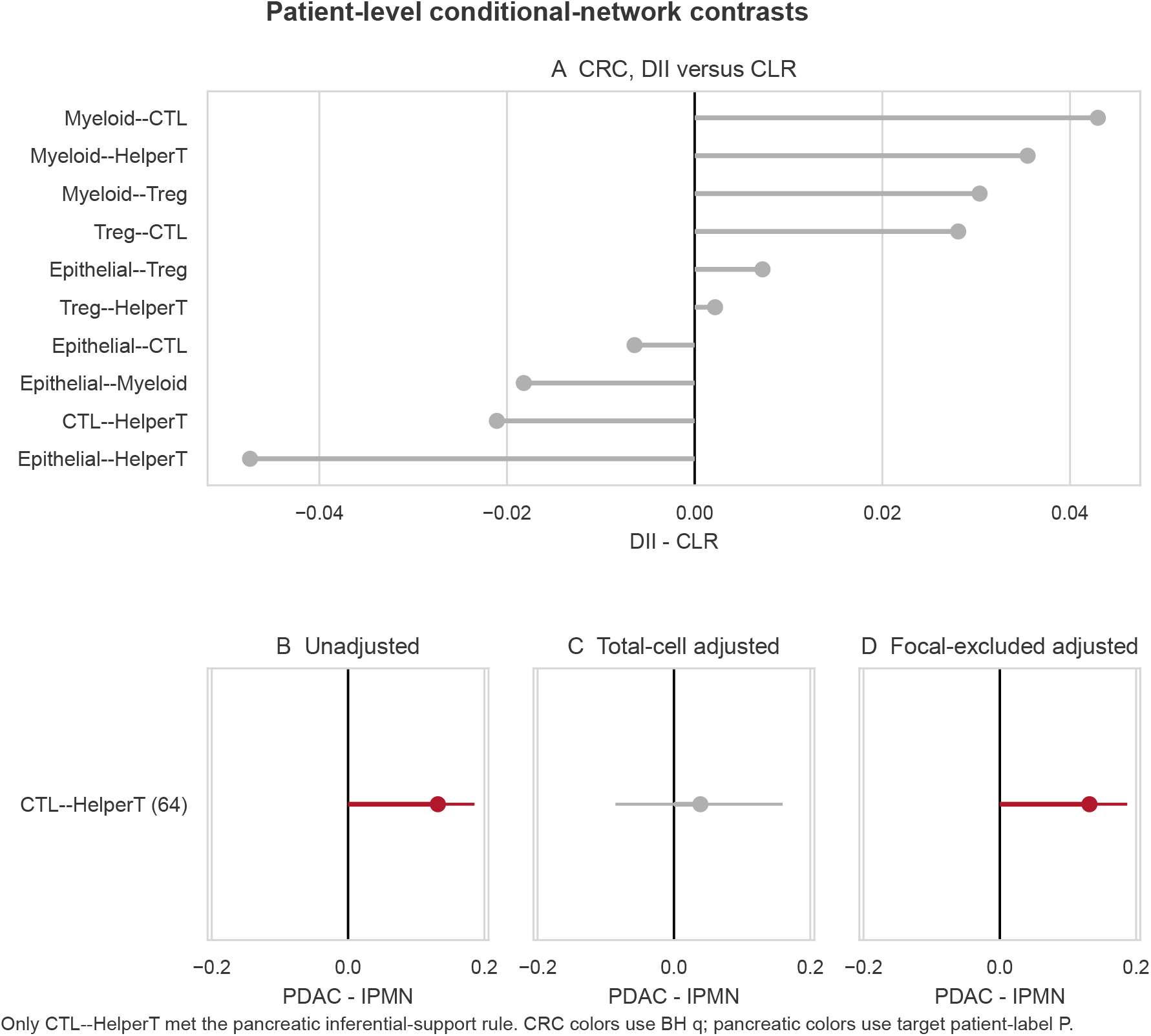
Patient-level differences in conditional-network edges. (A) CRC, DII minus CLR, averaged over the whole section. No global CRC edge contrast meets the FDR criterion, and the CRC result concerns the change of the Epithelial–Myeloid edge along the epithelial-density coordinate, which a section average cannot show. (B–D) Pancreatic CTL–HelperT contrast, PDAC minus IPMN, unadjusted (B), after total-cellularity adjustment (C), and after focal-excluded adjustment (D). Points are patient-level pathology contrasts and segments are 95% bootstrap intervals. Only CTL–HelperT is displayed for the pancreas because it was the only edge meeting the inferential-support rule (64 images with both densities varying); four computationally obtainable edges with 2–6 contributing images are omitted. Total-cell adjustment attenuates the contrast, whereas the focal-excluded adjustment retains it.

Schürch et al. (2020) described coordinated tumor, T-cell and macrophage neighborhoods in CLR and a more fragmented organization in DII. The present finding is consistent with that description and narrower. It locates the CLR–DII difference in the rate at which epithelial–myeloid coordination changes with local tumor density, and it shows that the difference is conditional and not marginal. This pattern may reflect a myeloid compartment that is displaced from, or excluded by, the densest epithelium more sharply in DII than in CLR, but the panel does not measure the function of these cells and the data are two-dimensional sections.

#### Conclusion

Within the estimands examined, the CRC signal is specific to conditional association and to its dependence on local epithelial density. To our knowledge, the rate at which epithelial–myeloid coordination changes along a within-section density coordinate has not previously been reported as the quantity that separates CLR from DII.

### 4.4 Colorectal cancer: recurrent neighborhoods and their change along the coordinate

K-means selected four CRC profiles, Low-density, Epithelial-rich, Treg-rich and CTL-rich, and three pancreatic profiles, APC-rich, Epithelial-rich and a T-cell-rich profile in which both CTL and helper T densities are elevated (standardized centers 1.62 and 1.69). The pancreatic silhouette was 0.777 at *K* = 3 against 0.740 to 0.505 for *K* = 4 to *K* = 7 (Figure 4). In the hierarchical patient/image bootstrap, the minimum proportion of replicates with matched-center correlation at least 0.8 was 0.960 across CRC profiles, so the profiles are properties of the cohort and not of one draw of patients. Along the epithelial-density coordinate, the prevalence of the CRC Epithelial-rich profile rose more rapidly from Q1 to Q4 in DII than in CLR, with a slope difference of 0.0571 per quartile (*q* = 0.0401), and the Low-density profile fell more rapidly in DII, with a slope difference of *−*0.0361 (*q* = 0.0404).

**Figure 4:**
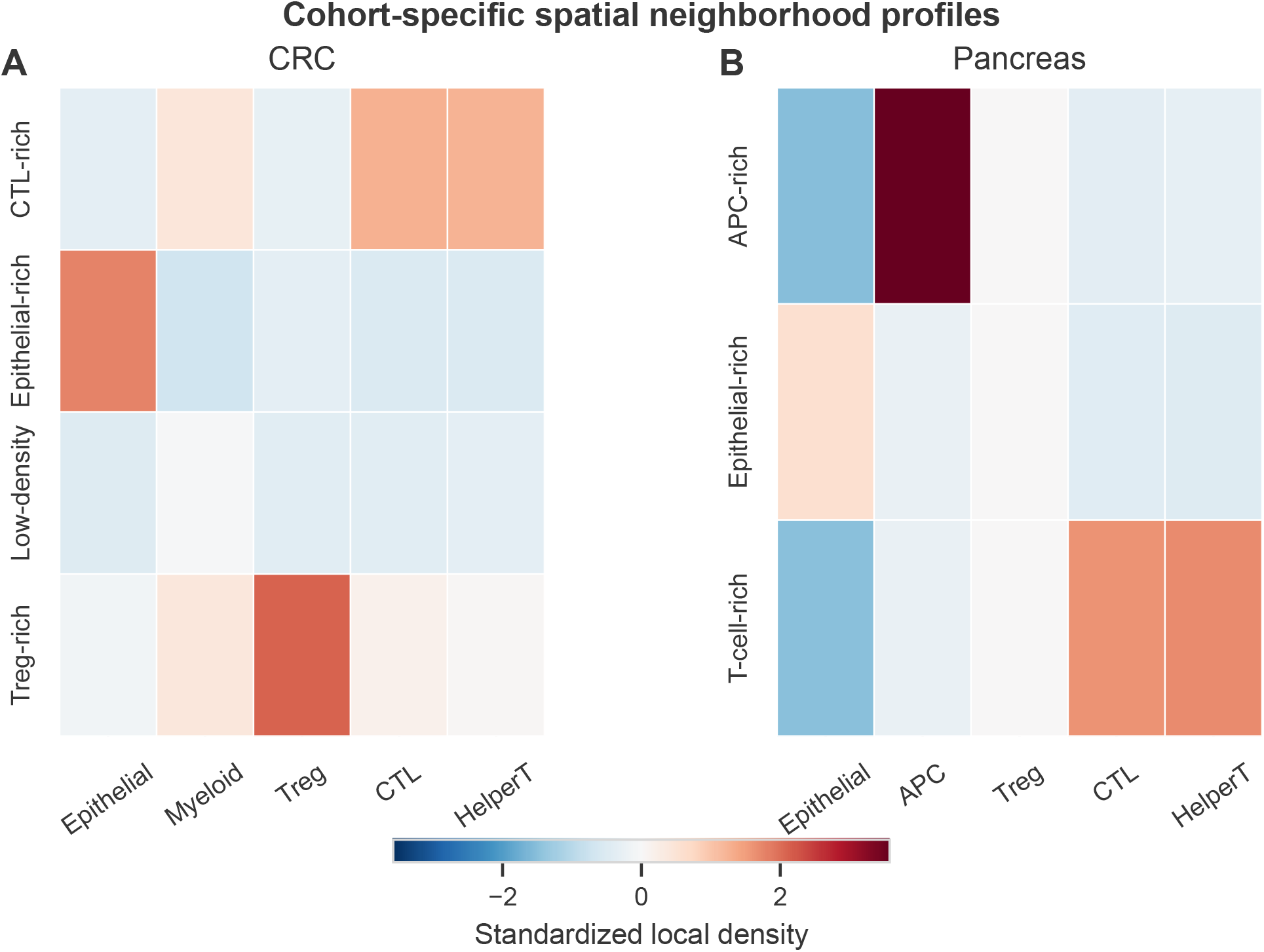
Estimated neighborhood centers by cohort. (A) CRC, four profiles. (B) Pancreas, three profiles. Coordinates are standardized phenotype densities. The pancreatic regulatory T-cell coordinate is constant and was excluded from profile naming and inference.

That recurrent neighborhoods distinguish CLR from DII corroborates the neighborhood analysis of Schürch et al. (2020), here with patients and not images as the resampling unit. That the epithelial-rich neighborhood accumulates faster along the within-section density coordinate in DII is, to our knowledge, not previously reported, and it is consistent with the steeper epithelial gradient in DII reported above. Note that these two statements are not independent, because a steeper epithelial gradient makes epithelial-rich bins more prevalent in Q4 by construction. The coordination result of Section 4.3 is not built in this way, since a partial correlation is invariant to the level of either density.

#### Conclusion

The CRC neighborhoods are reproducible under patient resampling, and their change along the coordinate is a descriptive companion to the conditional result, not independent evidence for it.

### 4.5 Pancreas: cytotoxic and helper T cells are closer together in PDAC

In the pancreas, the comparison rests on the one inferentially estimable pair, CTL–HelperT. We find that the two T-cell types are closer together in PDAC than in IPMN at every scale examined. Marginally, the differences in log_2_ enrichment were 0.463, 0.330 and 0.211 at *r* = 0.025, 0.05 and 0.10 (*q* = 0.0141, 0.0181 and 0.0229) across 33 IPMN and 45 PDAC patients. At the smallest radius, close CTL–helper-T pairs were about 1.4 times as frequent relative to random labeling in PDAC as in IPMN. The radii are fractions of a characteristic image length and not physical distances. Repeating the analysis under the earlier anisotropic normalization changed no conclusion (differences 0.448, 0.357 and 0.194; *q* = 0.0085, 0.0085 and 0.0310), so the finding is not an artefact of how sections were rescaled. Because the enrichment is computed against a random-label null that preserves each phenotype’s count, it is not a restatement of higher T-cell abundance in PDAC.

Organized helper and cytotoxic T-cell neighborhoods have been associated with longer survival in PDAC (Arseneau et al., 2025; Damle et al., 2026), and the T-cell-rich neighborhood profile of Section 4.4 places both densities together. The present finding corroborates that the two T-cell types are spatially associated in invasive disease and extends it to a precursor comparison, with the proximity greater in PDAC than in IPMN across three scales. This proximity may indicate that helper and cytotoxic T cells are recruited into shared niches as invasive disease develops, but the panel does not say whether the cells are in contact or functionally engaged.

#### Conclusion

Greater marginal CTL–HelperT proximity in PDAC than in IPMN is the pancreatic result that holds under every check we applied.

### 4.6 Pancreas: the conditional CTL–HelperT contrast depends on the cellularity nuisance

Conditionally, the answer depended on how local cellularity was represented (Figure 3B–D). Before adjustment, the patient-level CTL–HelperT partial correlation was *−*0.0778 in IPMN and +0.0540 in PDAC, a difference of +0.1318 across 66 IPMN and 55 PDAC patients (*|t|* = 4.80; target patient-label *P* = 0.001). The conservative five-edge sensitivity gave familywise *P* = 0.001 and BH *q* = 2.3 *×* 10*^−^*^5^. After residualizing each density on local total cellularity, the same contrast was +0.0388 and not distinguishable from zero (*|t|* = 0.61; target patient-label *P* = 0.520; five-edge familywise *P* = 0.960, BH *q* = 0.542). The focal-excluded nuisance did not reproduce this attenuation. At *λ* = 0.15, residualizing on *n*_other_ gave a PDAC-minus-IPMN difference of +0.1315 (95% bootstrap CI, 0.0817 to 0.1869; permutation *P* = 0.001).

This divergence arises because *n*_other_ barely varies in the 64 images where both T-cell densities vary (Table 1 gives the image counts). Among their 3,097 retained bins, its median was 0 (IQR, 0–0), 76.6% of bins had *n*_other_ = 0, and its correlation with total cellularity was 0.276. In other words, in these cores the T cells are nearly the only cells, so total cellularity and the focal counts are almost the same variable, and removing one removes the other. The divergence held at all five ridge penalties: total-cell-adjusted contrasts were weak and nonsignificant, and focal-excluded contrasts retained the unadjusted value (Supplementary Material). This is the behavior that abundance-sensitive spatial statistics have been shown to produce (Siddiqui et al., 2026; Soupir et al., 2025), observed here for a conditional and not a marginal estimand. It is new in one respect. The adjustment that removes the contrast and the adjustment that keeps it are both defensible, so the statement the data support is conditional on the nuisance definition.

#### Conclusion

Neither summary alone would have produced these statements. The marginal statistic cannot reveal the cellularity dependence, and the unadjusted network alone would have reported an association difference that does not survive total-cellularity adjustment. What the data support is that the two T-cell types are closer together in PDAC over the scales examined, and that whether their conditional association differs beyond cellularity depends on whether the nuisance burden includes the focal counts.

### 4.7 Sensitivity to spatial scale and to the network estimator

Disease-contrast edge rankings from the primary CRC grid had Spearman correlations of 0.842 and 0.867 with the coarser and finer grids, and edge signs agreed for 70% and 100% of CRC edges, respectively. The bandwidth refits of Section 4.3 and the neighborhood bootstrap of Section 4.4 assessed zoning and cluster sensitivity separately.

Figure 5 summarizes the comparison of the three network estimators in the 16-image CRC subset. The median within-image Spearman correlation between GP-GHS coefficient fields and localized ridge fields was 0.714 in CLR and 0.749 in DII (Figure 5A). GP-GHS and ridge assigned the same direction to 8 of 10 CRC pathology contrasts, and the Spearman correlation between their absolute contrast ranks was 0.855 (*P* = 0.00164). Agreement with ISPAT was weaker on both summaries (Figure 5B), which is the expected consequence of ISPAT averaging within zones while ridge and GP-GHS average over the section. All 16 selected CRC ISPAT fits completed. The methods condition on different variables and average over different scales, so this is concordance for selected motifs and not interchangeability. The pancreatic secondary-estimator comparison was performed on the superseded representation and is not carried forward. It supports no pancreatic conclusion here.

**Figure 5:**
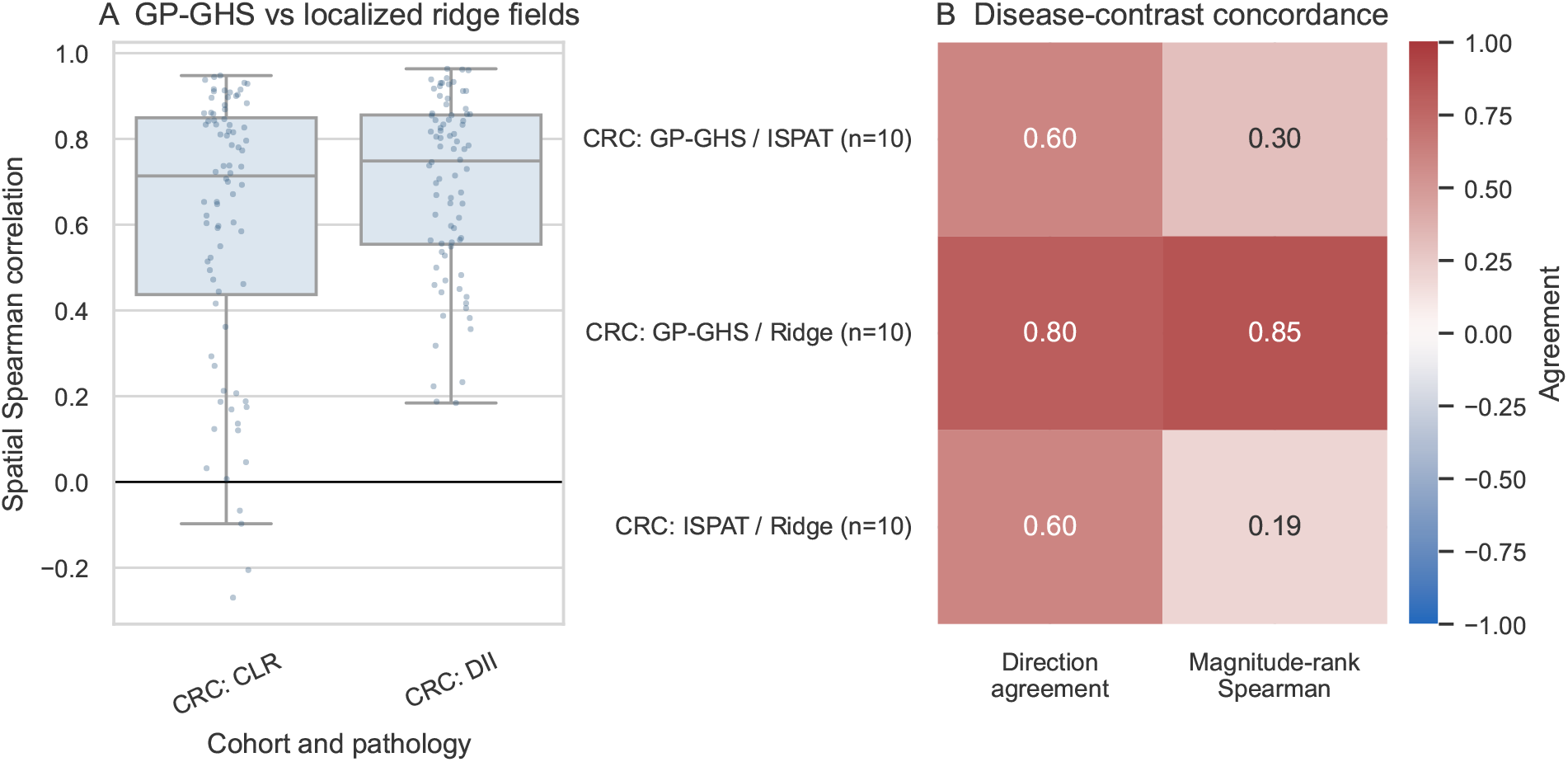
Comparison of ridge, ISPAT and GP-GHS estimates in the 16-image CRC subset. (A) Within-image spatial Spearman correlation between GP-GHS coefficient fields and localized ridge fields, by pathology. (B) Agreement in pathology-contrast direction and in absolute-magnitude rank for each pair of methods over the ten estimable edges.

### 4.8 Patient-level information carried by each spatial representation

We asked how much patient-level information about pathology each spatial representation carries when scaling, neighborhood-number selection, K-means fitting, held-out assignment, imputation and regression are all performed within each training fold. Figure 6 reports the leave-one-patient-out results. In CRC, neighborhood features gave the highest discrimination, AUC 0.794 (95% bootstrap CI, 0.608–0.948; *P* = 0.002), with Brier score 0.186 and calibration slope 0.785. Composition gave 0.742 (0.565–0.895; *P* = 0.011) and network features 0.690 (0.497–0.846; *P* = 0.029). Every one of the 35 folds selected *K* = 4. The combined model had lower AUC (0.719; 0.536–0.886) and a calibration slope of 0.340. Each training set held 34 patients while the combined model supplied 34 candidate predictors plus an intercept, so the ridge-edge features increased the parameter budget without adding stable signal. The *L*_2_ penalty permits estimation at this information ratio but cannot create incremental information or ensure calibration.

**Figure 6:**
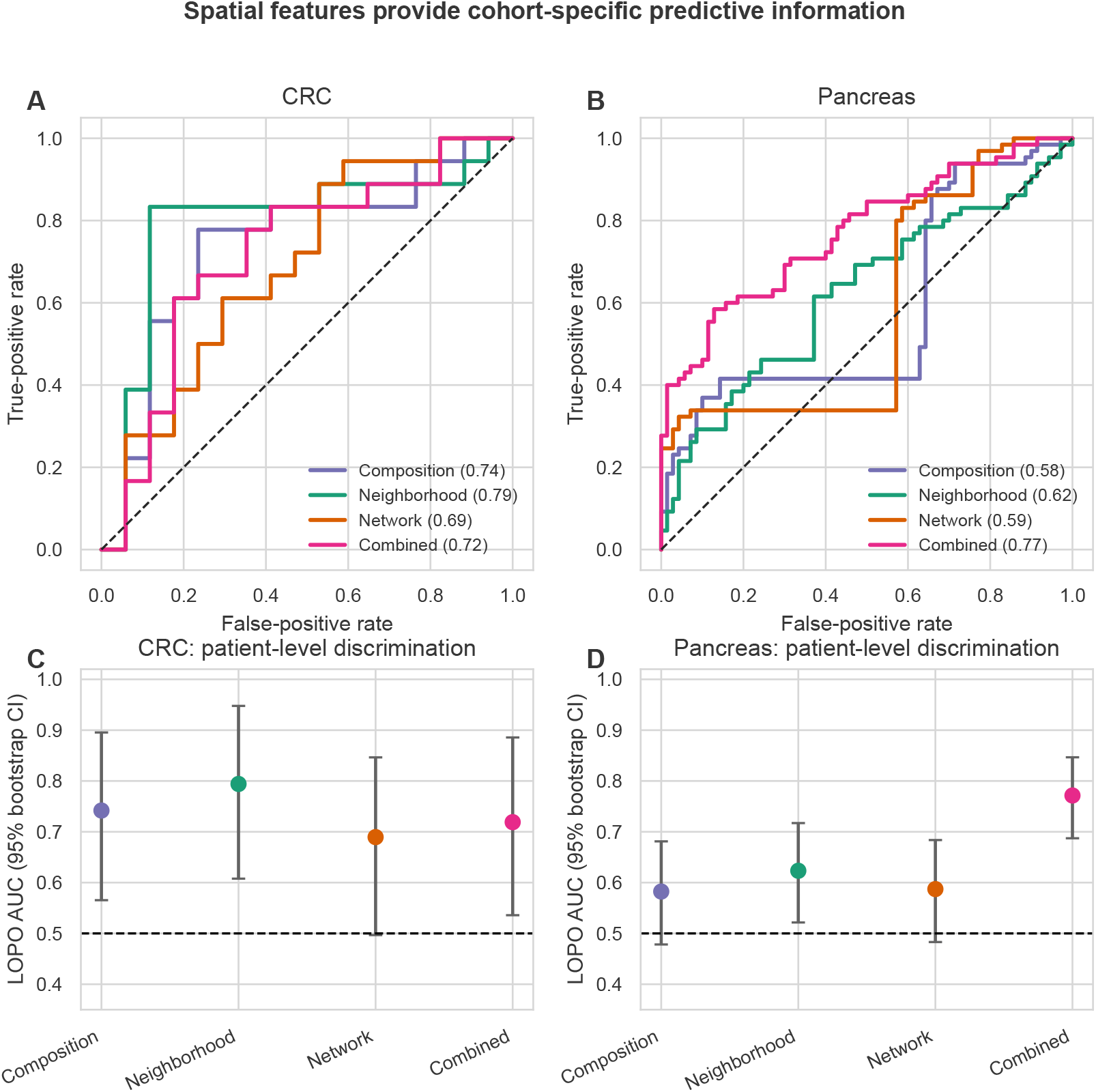
Leave-one-patient-out pathology discrimination. (A, B) Receiver operating characteristic curves for the four feature sets in CRC and in the pancreas, with AUC in the legend. (C, D) AUC estimates with stratified patient-bootstrap 95% intervals. Scaling, neighborhood-number selection, K-means fitting, imputation and regression were performed separately in each training fold.

In the pancreas, combined features gave AUC 0.771 (95% bootstrap CI, 0.687–0.847; *P* = 0.001) across 135 patients, against 0.623 for neighborhood features (0.522–0.717; *P* = 0.010), 0.587 for network features (0.483–0.684; *P* = 0.037) and 0.582 for composition (0.478–0.681; *P* = 0.049). All 135 folds selected *K* = 3. In summary, the representation that carries the most patient-level information differs by cohort. Recurrent neighborhoods carry it in CRC, where four images per patient make prevalences stable, and the combination carries it in the pancreas, where single cores make any one representation noisy and the four together are complementary.

These intervals and permutation values characterize uncertainty in the realized out-of-fold scores. They do not include variability from repeating the feature-learning pipeline, so feature construction, and not the uncertainty assessment, is what is fully nested. The models were not evaluated in an external cohort and include no clinical covariates, so they are internal estimates of pathology discrimination.

## 5 Discussion

We have asked how epithelial and immune cells are organized in the tumor microenvironment of two diseases, and we have answered with a patient-level analysis that keeps marginal proximity, recurrent local composition and conditional coordination as separate quantities. Applied to 35 CRC patients with 140 images and 141 pancreatic patients with 207 images, the analysis returned cohort-specific answers that differ in which quantity carries the signal. In CRC, the conditional epithelial–myeloid association changed along a within-section epithelial-density coordinate about four times faster in DII than in CLR, with no accompanying difference in proximity. In the pancreas, cytotoxic and helper T cells were more proximal in PDAC than in IPMN at every scale examined, and their unadjusted conditional association differed between the two diseases. However, that conditional difference disappeared once each density was residualized on local total cellularity and reappeared when the focal counts were excluded from the nuisance. The two cohorts are parallel applications of one standard and not replications of one biological result.

For the study of the tumor microenvironment, the findings carry two messages. The first is about colorectal cancer. Schürch et al. (2020) described coordinated tumor, T-cell and macrophage neighborhoods in CLR and a more fragmented organization in DII, and CLR is characterized by organized peritumoral lymphoid aggregates maturing toward tertiary lymphoid structures (Graham and Appelman, 1990). Our estimate is narrower than a neighborhood description. It is a rate of change along an epithelial-density coordinate, it was concordant when slopes were computed patient by patient and when the smoothing bandwidth was varied, and the edge kept its sign in both groups after cellularity adjustment. What the analysis adds to the neighborhood description is the estimand: a conditional association, at the patient level, as a function of local tumor density, which neither a proximity statistic nor a neighborhood prevalence can express. Because the CRC myeloid compartment combines CD68^+^ and CD163^+^ macrophages, the result concerns a myeloid density field and does not measure antigen presentation. The second message is about the pancreas. The robust conclusion is greater marginal CTL–HelperT proximity in PDAC, which is consistent with the organized T-cell neighborhoods that have been associated with survival in this disease (Arseneau et al., 2025; Damle et al., 2026). The conditional interpretation is sensitive to how cellularity is represented, and a single adjusted analysis would have hidden that sensitivity. We therefore do not claim that the conditional difference is uniquely explained by cellularity. This sensitivity may be common in tissue-microarray cohorts, where small cores are often dominated by one compartment, and it can be detected only when more than one nuisance definition is tried.

A more general message follows for how spatial organization is reported. Proximity and conditional coordination are different questions, and in these two cohorts they gave different answers in each direction: coordination without proximity in CRC, and proximity with a cellularity-dependent coordination in the pancreas. Cellularity adjustment is an estimand-defining choice, which is the addition this paper makes to the behavior that abundance-sensitive spatial statistics have been shown to produce (Siddiqui et al., 2026; Soupir et al., 2025). We suggest that studies of the tumor microenvironment report which of the three quantities a claim rests on, and report a conditional claim together with the nuisance definition it depends on.

A check that the two compartments are separable and supported had to precede both analyses. A conditional edge is interpretable only if its two density fields come from disjoint sets of physical cells, and it is estimable at the patient level only if enough images carry both fields. All ten CRC pairs met both requirements. In the pancreas the exclusive annotation made every pair separable, but only CTL–HelperT met the support rule, because epithelial and immune compartments almost never share a core (5 of 207 images contain at least three epithelial cells and at least three CTLs). This is a sampling property of a tissue-microarray cohort and not a limitation of any estimator, and it is why we report the pancreatic epithelial–immune quantities as non-estimable and not as null findings. We suggest that such a check belongs before, and not after, conditional network estimation in multiplex-imaging studies.

The prediction analysis compared how much patient-level information each spatial representation carries. Neighborhood features gave the highest AUC in CRC and combined features in the pancreas, with the neighborhood number constant across all folds in both cohorts. In CRC, adding network features raised the candidate set to 34 predictors plus an intercept for 34 training patients per fold, and AUC fell with a calibration slope well below one. Penalization made the fit computable but did not make weak added features informative. These are internally validated estimates of pathology discrimination, and we describe them as spatial discriminative signatures. External validation, clinical covariates and batch information would be required for any diagnostic use.

Two inferential choices deserve comment. First, a robust sandwich variance with 35 patient clusters remains a large-sample approximation, and our ordered-GEE simulation gave null interval coverage below nominal. We therefore treat patient-label randomization as the primary inferential statement for the CRC interaction and the clustered-GEE Wald test as a secondary model-based summary (Mancl and DeRouen, 2001). Second, the permutation statistic is studentized, so edges with different variances are compared on one scale. The CRC familywise value uses the per-permutation maximum across the ten estimable edges, whereas for the pancreas the target-edge test is primary and the maximum across the five obtainable edges is a conservative sensitivity analysis only.

Several limitations follow from the data. Both studies sampled two-dimensional sections, so apparent adjacency depends on the sectioning plane. Volumetric multiplexed imaging removes this dependence, and zone-specific conditional networks have recently been estimated on such data (Bhadury and Rao, 2026). IPMN and PDAC patients are independent groups, so nothing here estimates within-patient progression. No pixel-to-micron calibration was available, so proximity radii are fractions of a characteristic image length. We used an isotropic rescaling that preserves geometry, and the pancreatic conclusion was unchanged under the alternative. The pancreatic regulatory T-cell annotation contains no positive calls, and the pancreatic secondary-estimator comparison was performed on a superseded representation and is not carried forward. We could not adjust for clinical covariates (outcome, treatment or acquisition batch). Finally, the panels do not measure T-cell state or function, so none of the associations reported here is given a mechanistic meaning.

The analysis can be extended in two directions that the present data cannot reach. A hierarchical version can let the cellularity nuisance and the network parameters vary by patient, replacing the two fixed nuisance definitions with a model in which the coefficient on burden is a patient-level random effect, so that how much of an edge is explained by burden is estimated and not fixed by definition. The separability check can also be applied prospectively to a tissue-microarray design, choosing cores so that the compartments of interest co-occur often enough for their conditional association to be estimable, with the support rule of Section 3.1 as the design target. Experiments measuring T-cell state or function, and analyses in independent cohorts, are the next step before any of the associations reported here is given a biological meaning.

## Author contributions

S.B. conceived the statistical methodology, implemented the computational framework, performed all analyses, generated all figures, and drafted the manuscript. T.L.F. contributed to the pancreatic cohort and critically reviewed the manuscript. A.R. contributed to study conception and critically reviewed the manuscript. All authors reviewed and approved the final version.

## Generative AI use

OpenAI GPT-5.2 (OpenAI, 2025) was used to refine manuscript language and tone, format BibTeX citations, improve LaTeX displays (including algorithms and tables), and assist with R analysis and figure-generation scripts. GPT-5.2 and Claude Sonnet 4.6 (Anthropic, 2026) were also used to annotate, streamline, and document the R package for public release on GitHub. The authors reviewed and revised all AI-assisted text, code, and displays, checked the resulting outputs, and retain full responsibility for the methods, results, and conclusions.

## Data and code availability

The reusable implementation is publicly available in the spatialMIFtools GitHub repository. Manuscript-specific analysis scripts, configuration, input checksums, derived result tables and run logs are maintained together and will be archived as a versioned release; the immutable release identifier and archival DOI will be added at journal submission. The CRC CODEX data are available through The Cancer Imaging Archive under DOI https://doi.org/10.7937/TCIA.2020.FQN0-0326. The pancreatic mIF data are available from the University of Michigan School of Medicine upon reasonable request and institutional approval, subject to patient-privacy protections; requests may be directed to Marina Pasca di Magliano. Raw governed data are not redistributed by this project.

## Ethics

The CRC specimens and data were collected under approvals from the Ethics Committee of the Canton of Bern (KEK 200/2014) and the Stanford University Institutional Review Board (HSR 48803), as reported by the source study (Schürch et al., 2020). The pancreatic specimens were collected under University of Michigan Institutional Review Board protocol HUM00045286. The source pancreatic study reports written informed consent before enrollment, conduct in accordance with the Declaration of Helsinki, and HIPAA-compliant deidentification (Bhadury et al., 2026b). The present work is a secondary analysis of those deidentified, previously collected data and involved no new participant recruitment or specimen collection.

## Supplementary Material

The Supplementary Material, provided as a separate file, contains the patient-label permutation distributions, the cellularity-definition and ridge-penalty sensitivity table, the coordinate-normalization comparison for cross-pair enrichment, and the graphical summaries of the simulation experiments.

